# Transcriptomic analysis of cells following decreased mitochondrial DNA-copy number reveals compensatory mechanisms in mitochondrial DNA replication and cellular energetics

**DOI:** 10.1101/2025.11.10.687605

**Authors:** Jiaqi Xie, Phyo W. Win, Charles Newcomb, Shaopeng Zeng, Christina A. Castellani, Dan E. Arking

## Abstract

Mitochondrial DNA copy number (mtDNA-CN) is a metric of mitochondrial function that has been associated with a variety of diseases including cardiovascular disease and all-cause mortality. To investigate genes and pathways affected by mtDNA-CN variation, we perturbed HEK 293T cells with ethidium bromide to deplete mtDNA. Using RNASeq and methylation microarrays, we evaluated transcriptomic and methylomic changes in treated cell lines. We observed an 8-fold decrease in mtDNA-CN and compensatory shifts in mitochondrial transcription to support mtDNA replication. Nuclear transcriptomic and methylomic analysis highlighted changes in metabolic pathways, including oxidative phosphorylation and canonical glycolysis. Longitudinal analyses revealed that the identified genes and pathways have different response timing, with nuclear response lagging behind mitochondrial response. These findings further elucidate the mechanisms behind mtDNA maintenance and responses to cellular energetics as well as mitochondrial-nuclear crosstalk dynamics.

## 1 Introduction

Among the organelles of the mammalian cell, only the mitochondrion maintains its own DNA, a circular genome measuring 16,569 base pairs long encoding 2 ribosomal RNAs, 14 transfer RNAs, and 13 protein subunits of the oxidative phosphorylation pathway.^1^ Although these genes only account for approximately 1% of all proteins localized to the mitochondria, maintaining mitochondrial DNA (mtDNA) integrity is essential for cellular and organismal health.^2^ Mitochondrial DNA-copy number (mtDNA-CN), as a metric of mtDNA abundance and mitochondrial function, has been associated with cardiovascular disease, neurodegenerative disorders, and all-cause mortality, among other disease states.^3–6^

Previous work has shown that blood mtDNA-CN is associated with altered gene expression, with neurodegenerative disease pathways highlighted by genes correlated with measured mtDNA-CN.^5^ A separate study demonstrated that individuals with lower mtDNA-CN also display differential methylation of the nuclear genome, and the differentially methylated CpG sites are enriched for associations with cardiovascular disease.^7^ The idea that mtDNA-CN variation could affect disease outcomes through altering nuclear methylation and gene expression is additionally supported by observations in *TFAM* knock-out cell lines demonstrating that cell signaling pathways are overrepresented by affected gene-CpG pairs.^8^

In this study, we aimed to identify nuclear encoded genes and pathways responsive to variations in mtDNA-CN. We exposed cell lines to ethidium bromide (EtBr), a chemical known to intercalate with and deplete mtDNA.^9,10^ First, we demonstrate that mtDNA-CN responds to EtBr in a dose dependent manner and identify changes in gene expression and nuclear DNA methylation associated with decreasing mtDNA-CN. Second, we show that cell lines subjected to EtBr treatment and recovery have distinct temporal patterns in mtDNA-CN and gene expression, establishing the timing at which each gene responds to mtDNA-CN variation. Third, we suggest that changes in the expression of the mitochondrial genome arise from prioritizing mtDNA replication over transcription. Finally, we observe that affected pathways tend to be made up of genes showing a delayed temporal response compared to the mtDNA-CN response, and that the cell compensates for decreased mitochondrial function by cutting down on energy expenditure, while allocating additional resources to oxidative phosphorylation and switching to alternative energy sources.

## 2 Methods

### 2.1 Culturing HEK293T cells

Human embryonic kidney 293T cells (ATCC CRL-3216) were plated at a density of 200,000 cells/well within 6-well plates in high glucose DMEM medium (Gibco 11965118) supplemented with 10% fetal bovine serum and 1% penicillin-streptomycin. Cells were placed in a 37°C, 5% carbon dioxide incubator for 24 hours to allow the cells to adhere to the plates.

### 2.2 Ethidium Bromide dosage experiment

For chemical treatment, medium was removed and replaced with fresh DMEM supplemented with varying concentrations of ethidium bromide (EtBr). Seven different concentrations of EtBr (0, 25, 50, 75, 100, 125, 150 ng/mL) were used, with 0 ng/mL used as control. Cells were then grown for 48 hours following treatment before harvesting. All treatment conditions were performed in triplicate (N=3).

### 2.3 Treatment and recovery experiment

The effect of treatment and removal of EtBr was assessed by growing 6 wells of cells in media with 150 ng/mL EtBr and 6 wells of cells in media with no EtBr. One well for each group was harvested at times 0, 24, and 48 hours. At 48 hours, the medium was removed and replaced with fresh DMEM without EtBr for all remaining wells, which were harvested at 72, 120, and 144 hours. Experiments were performed in triplicate (N=3).

### 2.4 DNA extraction

Experimental cells were harvested, pelleted, and further lysed in RLT Plus Lysis Buffer (Qiagen 1053393). Phenol:chloroform:isoamyl (PCIA) alcohol (Invitrogen 15593031) was used to extract the DNA-containing solute with sodium acetate (pH 5.2, Quality Biologicals 351-035-721), 100% ethanol (ETOH), and glycogen. DNA was precipitated overnight with additional ETOH washes. The resultant DNA samples were checked for quality using NanoDrop, and all samples had 260/280 within 1.8 – 2, showing good quality. Samples were stored at -20°C until further use.

### 2.5 Monochrome multiplex qPCR of mtDNA-copy number

To estimate mtDNA-CN, we adapted the protocol described by Hsieh et al. (2018).^11^ DNA samples were quantified using a Qubit 2.0 Fluorometer and Broad Range dsDNA assay (Invitrogen Q32850). Samples were diluted to 10 ng/uL using molecular grade water for use in the qPCR reaction. mtDNA-CN was assayed by comparing relative cycle threshold differences between the nuclear-encoded albumin gene (forward primer: 5’ – CGG CGG CGG GCG GCG CGG GCT GGG CGG AAA TGC TGC ACA GAA TCC TTG – 3’; reverse primer: 5’ – GCC CGG CCC GCC GCG CCC GTC CCG CCG GAA AAG CAT GGT CGC CTG TT – 3’) and the mitochondrial D-Loop (forward primer: 5’ – ACG CTC GAC ACA CAG CAC TTA AAC ACA TCT CTG C – 3’; reverse primer: 5’ – GCT CAG GTC ATA CAG TAT GGG AGT GRG AGG GRA AAA – 3’). DNA primers were purchased from IDT. Each qPCR reaction consisted of 10 uL total volume comprised of 5uL of 2x Quantifast SYBR Green Master Mix (Qiagen 204056), 0.25 uL of each albumin and D-Loop primer to a final concentration of 1 uM, 2 uL of 10 ng/uL, DNA stocks for 20 ng input, and 2 uL of molecular grade water. All qPCR reactions were run in triplicate. Reactions were run on a Viia7 Real-Time PCR System (Applied Biosystems) in a 384-well PCR plate (Applied Biosystems 4309849). Thermocycling conditions for the reactions were: 95°C incubation for 10 minutes, then 40 cycles of 94°C for 15 seconds, 62°C for 10 seconds, 74°C for 20 seconds (with data acquisition), 84°C for 10 seconds, and 88°C for 20 seconds (with data acquisition). Ramping speeds were set to 1.9°C/second at all stages. We determined raw mitochondrial DNA copy number for each sample by calculating the difference in cycle threshold (Ct) values between the nuclear- and mitochondrial-specific PCR products from the above primer sets. Each sample was measured 3 times, and measurements with D-Loop values >26 or delta Ct values <7 were filtered out. The average of the remaining replicate measurements for each sample was used as the estimate for mtDNA-CN.

### 2.6 Measuring nuclear DNA Methylation

Nuclear DNA methylation measures were performed using isolated DNA. Bisulfite conversion of nuclear DNA was performed using the EZ-96 DNA Methylation Kit (Deep Well Format) (Zymo Research D5004; Irvine, CA, USA) according to the manufacturer’s protocol. Following bisulfite conversion, DNA was hybridized to the Illumina Infinium Methylation EPIC Beadchip microarray at the Erasmus University Medical Center Rotterdam to determine nuclear DNA methylation profiles. Methylation data was processed using the ‘minfi’ R software package and poor-quality probes were removed if the probe had a detection p-value > 0.01.^12^ The data was normalized using functional normalization.^13^ Cross-reactive probes were identified and removed, as well as probes with known SNPs at the CpG site.^14^ Principal component analysis of methylation values was used to identify outliers. Specifically, samples were removed if they were 3 or more standard deviations from the mean in any of the first ten principal components.

### 2.7 Determining differential DNA methylation

Differentially methylated sites were determined using a mixed linear regression model with mtDNA-CN as the independent variable and methylation values as the dependent variable, as well as the inclusion of methylation chip as a random variable. A significance threshold of p-value < 0.05 after Holm–Bonferroni multiple testing correction was used to determine differentially methylated sites.^15^

### 2.8 RNA extraction

Total RNA was extracted using the RNeasy Plus Mini Kit (Qiagen 74134) according to manufacturer’s protocol. Samples were eluted from the column twice using 50 uL of molecular grade water. Samples were then stored at -80°C. Fifty-seven total samples were each sequenced twice. An internal control sample (a repeated negative control sample from the EtBr dose curve) was also included.

### 2.9 Quality control of RNA samples

RNA sample quality was assessed using both NanoDrop and Fragment Analyzer (Advanced Analytical Technologies). It is expected for high quality RNA samples to have 260/280 measurements of approximately 2.0 when measured on the NanoDrop, and the average 260/280 across all 112 samples was 2.03 ± 0.03. RQN scores from Fragment Analyzer for 111 of the 112 samples were 10.0, the highest possible measure and indicating high-quality RNA samples. One sample from cells grown in 100 ng/mL EtBr had a RQN score of 6.1; this sample was still included in library preparation since NanoDrop measures were still acceptable and subsequent analysis showed no issues with the sample.

### 2.10 cDNA library prep

cDNA libraries were prepared following the manufacturer’s protocols using the NEBNext Ultra II RNA Library Prep Kit for Illumina (NEB E7775) and the Poly(A) mRNA Magnetic Isolation kit (NEB E7490). The protocol was optimized for 200 bp RNA inserts. Briefly, 900 ng of each sample was used as input. Total RNA was washed, and mRNA was isolated and then fragmented. Double-stranded cDNA was synthesized, and adapters were added onto all samples. Unique dual index primer pairs were used to barcode each sample. Specifically, all indexes from ‘Plate 1’ (NEB E6440) and the first two columns from ‘Plate 2’ (NEB E6442) were used. Following manufacturer recommendations, samples were PCR-amplified for 8 cycles, cleaned via magnetic beads, and eluted in 1x TE Buffer, resulting in completed cDNA libraries.

### 2.11 Quality control of cDNA libraries

Library quality was assessed using the Fragment Analyzer and associated DNA kit. Initial QC showed a primer-dimer peak in most sample traces. To remove the dimers, samples were first diluted to 4 nM using the molarity determined by the Fragment Analyzer for the region excluding the primer dimer peaks (200 to 1000 bp region of the trace). 4 uL of each diluted library was then pooled together, and the pool was purified using a 1x magnetic bead clean up. The pool was then assessed for quality on a BioAnalyzer 2100 using the High Sensitivity DNA Assay (Agilent), and removal of the dimers was confirmed. The calculated molarity of the pooled samples after clean-up was 12.5 nM and average fragment size was 410 bp.

### 2.12 RNA sequencing

RNA sequencing was performed by the High Throughput Sequencing Center of the Genetic Resources Core Facility at Johns Hopkins University. The cleaned and pooled samples were submitted at 4 nM in 50 uL volumes. Sequencing was performed on the Illumina NovaSeq system using an S2 flowcell and v1.5 reagents. Quality of the RNAseq data was checked in-house using FASTQC, and all sequences had passing per sequence quality scores.

### 2.13 Statistical analysis and visualization

All statistical analysis was conducted with R statistical software v4.3.0 using RStudio.^16,17^ Data visualization and plots were generated with the aid of the following packages: ‘ggplot2’ (v3.5.1), ‘ggfortify’ (v0.4.17), ‘ggh4x’ (v0.3.0), ‘ggpubr’ (v0.6.0), ‘qqman’ (v0.1.9), ‘RColorBrewer’ (v1.1-3).^18–24^

### 2.14 Deriving normalized read counts from RNA sequencing data

We used Trimmomatic (v.0.39) to remove adaptor sequences and filter reads for Phred quality score >20 and read length >50 bp.^25^ We added ‘N’s to reads to ensure that paired reads were the same length. Paired reads were aligned with RUM (v2.0.5_06) to the GRCh38 (hg38) reference genome and transcriptome.^26^ We then counted the number of alignments mapped to each gene with HTSeq (v2.0.7) using a gene transfer format file from Ensembl (v.99). ^27,28^ We removed genes with median counts lower than 50 in the control samples.

We used principal component analysis (PCA) to calculate the first 10 principal components of the RNA counts and removed any samples with a principal component more than 4 standard deviations from the mean.^29^

Finally, to account for differences in library size we normalized read counts for each gene using Trimmed Mean of M values (TMM) as implemented by the ‘edgeR’ (v4.0.6) package in R.^30,31^

### 2.15 TWAS identifying genes associated with mtDNA-copy number

For each gene, we took measurements from the dosage experiments and log transformed read counts were regressed against mtDNA-CN using a linear mixed model, implemented in the ‘lmerTest’ (v3.1.3) package in R,^32^ to account for multiple measurements of each sample.

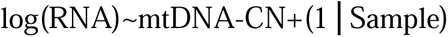

Significantly differentially expressed genes were identified as those with p-values <0.05 after Holm–Bonferroni multiple testing correction.^15^

### 2.16 Categorizing longitudinal behavior of genes and CpGs

Significantly differentially expressed genes were additionally categorized by their longitudinal behavior in the treatment and recovery experiments. Log fold-change (logFC) between the treated and control cells were calculated at hours 0, 24, 48, and 72. LogFC were scaled so that the mean logFC was 0 at hour 0 and 1 at hour 48. We used Bayesian posteriors to calculate the probability that a gene with given scaled logFC would follow one out of three longitudinal patterns: 1) log-linear behavior; 2) switch-like behavior; and 3) temporally delayed behavior. Expected scaled logFC for each of the patterns is described below. The prior likelihood was assumed to be equal for all three patterns. The posterior possibility was calculated using the likelihood of observing the scaled logFC at 24 and 72 hours assuming a normal distribution for measured logFC.

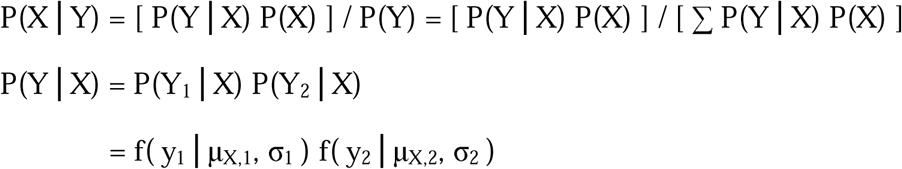

f is the normal probability density function for a given mean and standard deviation.

X∈{1,2,3}indicates longitudinal pattern, prior probability P(X)=1 ⁄ 3

μ_X,1_, μ_X,2_ are expected logFC observed at 24 and 72 hours.

Pattern 1: μ_1,1_ = 0.5, μ_1,2_ = 0.5

Pattern 2: μ_2,1_ = 1, μ_2,2_ = 0

Pattern 3: μ_1,1_ = 0.25, μ_1,2_ = 1

Y is Observed data, Y_1_ and Y_2_ are measurements at 24 hours and 72 hours respectively

Mean and standard deviation of Y_i_

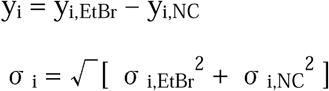

A gene was assigned to a particular pattern if the posterior possibility for that pattern is at least twice that of any other pattern, otherwise the gene was labeled ‘ambiguous’. Sensitivity analyses were conducted by relaxing the cutoff to assigning the pattern with the greatest posterior probability or tightening the cutoff by requiring posterior probability to be at least 4 times that of other patterns.

The same categorization framework was used for methylation sites where a methylation CpG was assigned to one of the longitudinal patterns based on methylation level and directionality. Additionally, significant differentially methylated CpGs were mapped to nearby differentially expressed genes with a delayed pattern within a range of 1Mbp to identify CpG-Gene pairs.

### 2.17 Enrichment analysis

We conducted Gene Ontology (GO) enrichment analysis, also called overrepresentation analysis, on the set of significantly upregulated and downregulated genes, as well as on the set of differentially expressed genes near differentially methylated CpGs (CpG-Gene pairs).^33–35^ Given that EtBr treatment decreases mtDNA-CN, upregulated genes were defined as those with a negative estimate and significant p-value after Holm–Bonferroni multiple testing correction, while downregulated genes were those with a positive estimate and significant p-value after Holm–Bonferroni multiple testing correction. Additionally, we conducted enrichment analysis for up and downregulated genes assigned to each of the three longitudinal patterns. Enrichment analysis was performed using the ‘clusterProfiler’ (v4.10.1) and ‘limma’ (v3.58.1) packages in R.^36,37^ We identified GO biological processes as over-represented by differentially expressed genes if they had p-value < 0.05 after Benjamini-Hochberg multiple testing correction.^38^ This more lenient Benjamini-Hochberg multiple testing correction was used instead of Bonferroni-Holm for enrichment analysis because GO terms are heavily correlated with one another and violate the assumption of independent tests. Similar significant terms were grouped together using the ‘rrvgo’ (v1.14.2) package (Supplementary Fig 1).^39^

## 3 Results

### 3.1 mtDNA-CN decreased with increasing doses of EtBr

To identify genes associated with variation in mtDNA-CN, we generated a gradient in mtDNA-CN. Cells were grown in different doses of EtBr, from 0 ng/ml to 150 ng/ml, for 48 hours. With each 25 ng/ml increase in EtBr dosage, we saw mtDNA-CN decreasing significantly (p < 0.05) until 125 ng/ml, with cells grown in 125 ng/ml and 150 ng/ml EtBr showing similar levels of mtDNA-CN (Fig. 1).

**Fig. 1.**
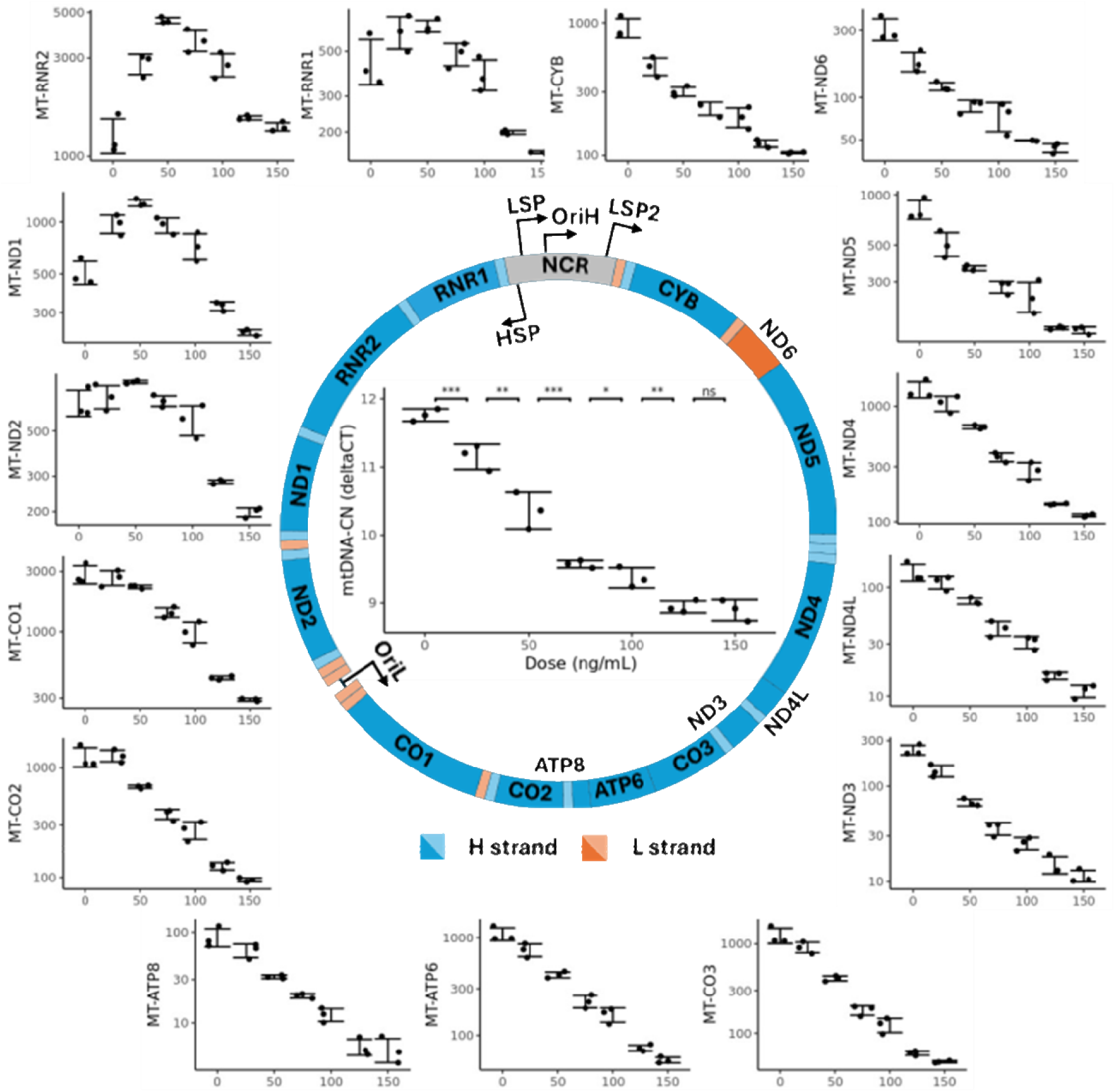
Mitochondrial DNA and RNA abundance decrease due to EtBr treatment. Circular diagram shows the human mitochondrial genome, surrounded by plots showing changes in mtDNA-CN and mtRNA counts for cells grown in different dosages of EtBr (N=3). Mitochondrial DNA-copy number is in the center, and mtRNA are arranged by the position of the gene. Error bars are +/-1SD.

### 3.2 Mitochondrial DNA encoded genes close to the heavy strand promoter (HSP) showed nonlinear decreases in mapped reads with increasing EtBr dosage

Previous work has demonstrated that the levels of mitochondrial RNA (mtRNA) are associated with mtDNA-CN.^5^ Given the high turnover and short half-lives of mtRNA, we expected a decrease in mtDNA-CN to be reflected by a decrease in mtRNA.^40^ We examined whether mapped reads for specific genes have varied responses to EtBr treatment. Since mature tRNAs were excluded by fragment size selection during RNA-Seq library preparation, we only analyzed reads mapped to the protein coding genes and rRNAs. We observed that 11 protein coding genes showed a log-linear decrease in gene expression with increasing EtBr dosage, while the 2 rRNAs (*MT-RNR1*, *MT-RNR2*) and 2 additional protein coding genes (*MT-ND1*, *MT-ND2*) exhibited nonlinear responses. Read counts for these four genes initially trended upwards, peaked at 50 ng/mL EtBr, and trended downwards as EtBr dosage increased further (Fig. 1). Nonlinear responses from genes close to the HSP suggest that transcriptional initiation has increased following EtBr treatment.

### 3.3 Fold changes in mtRNA mapped reads were correlated with the distance between each gene and its closest promoter

To further examine the differential response of mtRNAs to EtBr treatment, we calculated logFC of mapped reads for each mtDNA gene and plotted them against their position on the mitochondrial genome. We observed that logFC is mostly consistent at all positions when EtBr dosages were low, but a U-shaped pattern becomes more prominent as EtBr dosage increased with prominent bowing in the center (Fig. 2A). Instead, when we plotted logFC for cells grown in 150 ng/mL EtBr against the distance from each gene to the closest promoter, a significant linear pattern emerges (Fig. 2B, P = 4.02×10^-3^). The symmetry of the response suggests that sense and antisense reads originating from both transcription sites were being captured, with processivity decreasing as distance increases from the promoter, acting together with decreased mtDNA-CN to lower mtRNA expression.

**Fig. 2.**
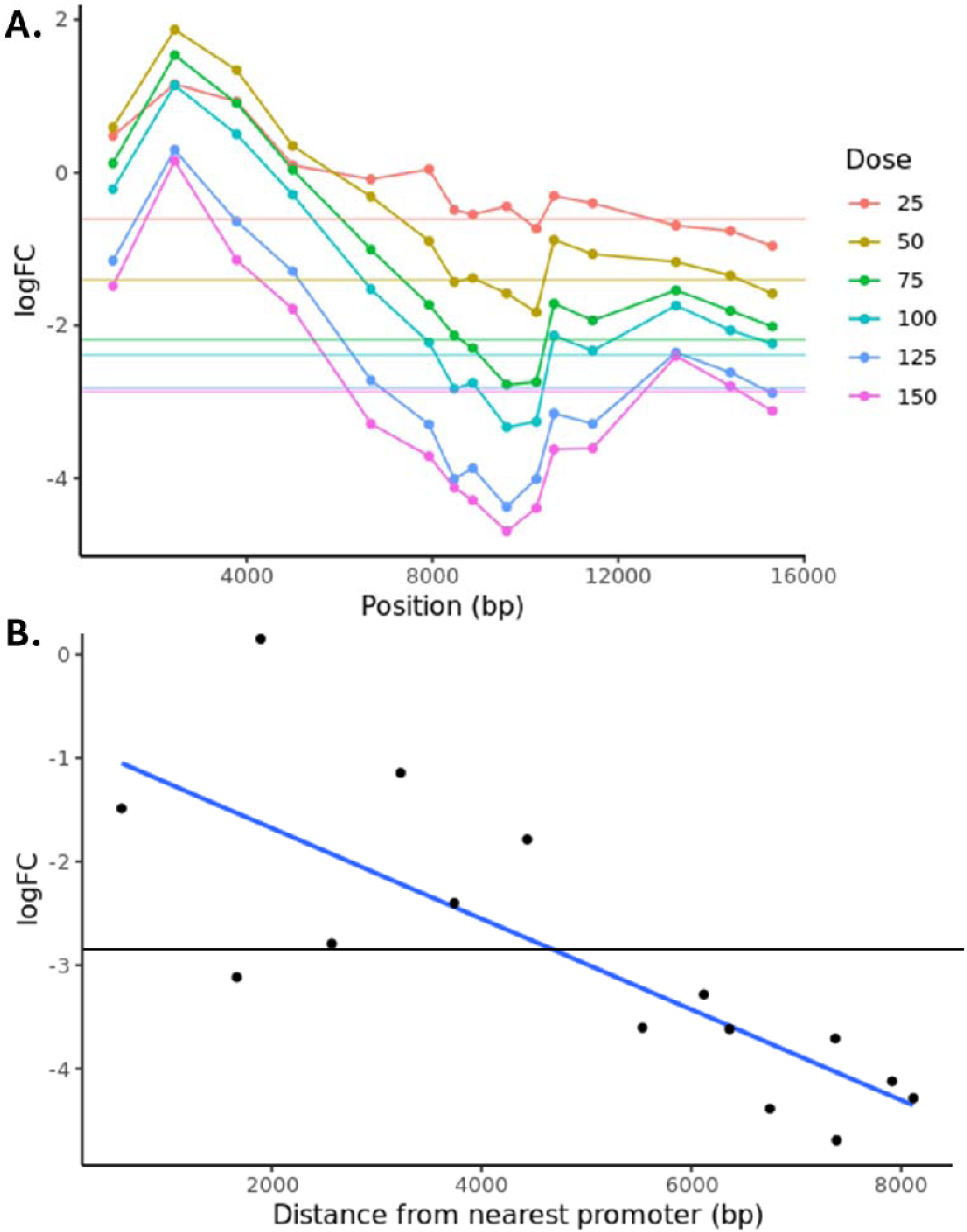
mtRNA read counts exhibit non-uniform decrease after EtBr treatment. **A.** The logFC of individual mtRNAs, as shown by the points, shows positionally dependent deviation from logFC of mtDNA-CN in EtBr treated cells, as shown by the horizontal lines. **B.** In 150 ng/mL EtBr cells, mtRNA logFC shows a linear pattern when plotted against the distance from each mtRNA to the closest promoter. Horizontal line denotes logFC in mtDNA-CN.

### 3.4 Transcriptome-wide association study (TWAS) between mtDNA-CN and nuclear encoded genes

Using the gradient in mtDNA-CN, we tested if nuclear encoded genes were affected by variation in mtDNA-CN by regressing nuclear gene expression against mtDNA-CN. Of the 11,570 highly expressed nuclear encoded genes, 390 were significantly differentially expressed (DE) after multiple-test correction, with 199 genes upregulated and 191 downregulated. Note that because EtBr treatment decreases mtDNA-CN, upregulated genes have negative coefficients while down regulated genes have positive coefficients. Top downregulated genes included *TCTN1, ANAPC1*, *CEBPG*, *PRSS23*, and *STC2*. Top upregulated genes included *TBCD*, *G6PD*, *IDH3G*, *GET3*, and *RFLNB* (Supplementary Table 1).

Using overrepresentation analysis, we found that multiple biological pathways, as annotated by Gene Ontology (GO), were affected by the decrease in mtDNA-CN, with 18 GO terms upregulated and 10 terms downregulated (Fig. 3). We grouped enriched GO terms by their similarity. Upregulated GO terms included amino acid metabolic process (GO: 0006520, adj.p = 1.54×10^-9^), tRNA aminoacylation (GO:0043039, adj.p = 5.31×10^-8^), and amino acid import across plasma membrane (GO:0089718, adj.p = 1.92×10^-3^). Downregulated terms included cholesterol biosynthetic process (GO:0006695, adj.p = 7.35×10^-4^) and mRNA processing (GO:0006397, adj.p = 0.0231). Contrary to expectations, nuclear encoded mitochondrial processes were not significantly enriched.

**Fig. 3.**
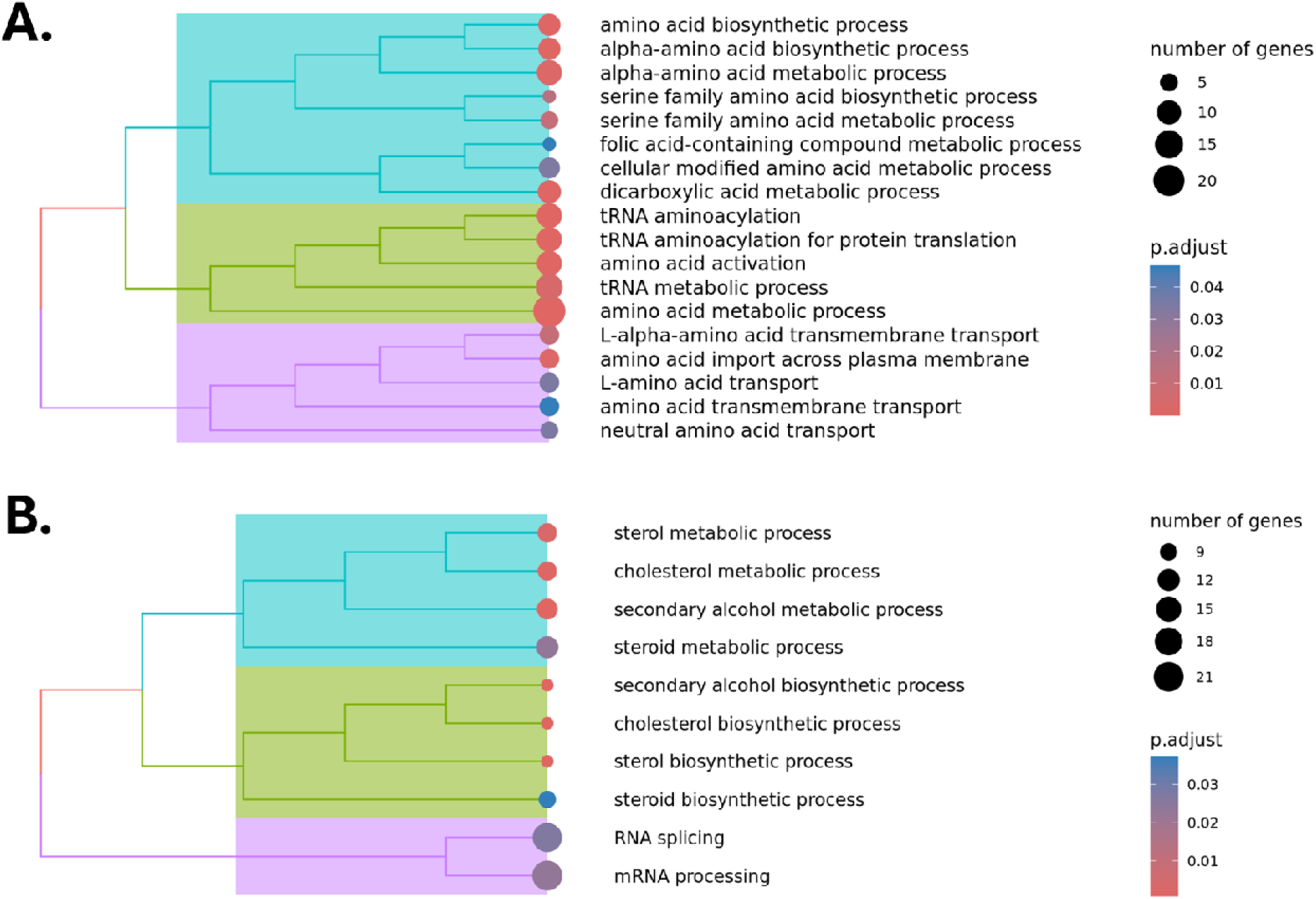
GO Pathways enriched by differentially expressed genes. Overrepresentation analysis was conducted on A) upregulated genes, and B) downregulated genes.

### 3.5 Nuclear encoded mitochondrial genes for OXPHOS were differentially expressed in the presence of EtBr

We more closely examined the nuclear genes involved in mitochondrial maintenance and the oxidative phosphorylation (OXPHOS) pathway, as we had expected these pathways to be affected by changes in mtDNA/mtRNA abundance induced by EtBr treatment. We therefore focused on genes assigned to these pathways via protein localization and literature curation in MitoCarta3.0.^2^ For genes responsible for mtDNA maintenance, mtRNA metabolism, and translation, we found equal numbers of significantly upregulated and downregulated genes in each pathway, suggesting that increased transcription initiation and mtDNA synthesis within the mitochondria is not reflected by changes in nuclear gene expression (Table 1).

**Table 1.**
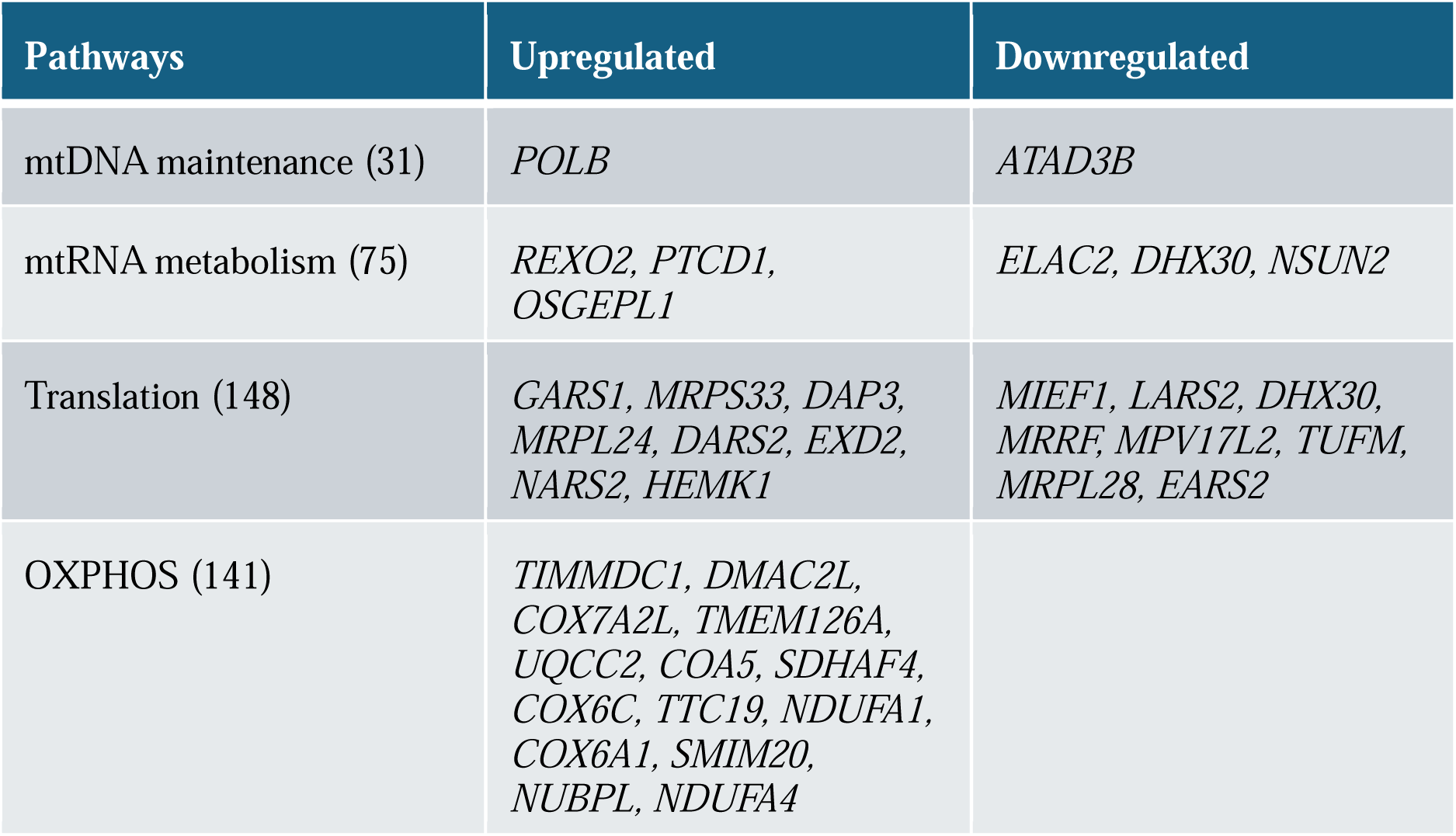
Differentially expressed (adj.p < 0.05) genes in mitochondrial pathways. Number in parentheses indicate total number of genes in each pathway.

In contrast, of the 141 genes involved in oxidative phosphorylation, 14 were significantly upregulated while none were significantly downregulated. While mitochondrial ATP synthesis coupled electron transport (GO:0042775) was not significantly overrepresented by the upregulated genes, the skewed distribution of the differentially expressed genes suggests that the pathway was upregulated by decreased mtDNA-CN.

### 3.6 Glycolysis genes exhibited nonlinear response to increased EtBr dosage

Prior experiments in EtBr treated cells showed that depletion of mtDNA decreased ATP synthesis via the electron transport chain, and glycolysis increased to compensate.^41^ We examined genes involved in canonical glycolysis expecting to see a similar response. Instead, we observed that expression of glycolytic genes decreased with increasing concentration of EtBr. Some genes in the canonical glycolytic pathway (*ALDOA, TPI1, ENO1, PKM*) showed a fishhook-like nonlinear pattern, with gene expression reaching a minimum at 100 ng/mL EtBr before reversing direction and trending upwards as EtBr concentration increased to 150 ng/mL (Fig. 4). Nonlinear down regulation of canonical glycolysis genes suggests that the pathway is not solely regulated by the cell’s energy demands.

**Fig. 4.**
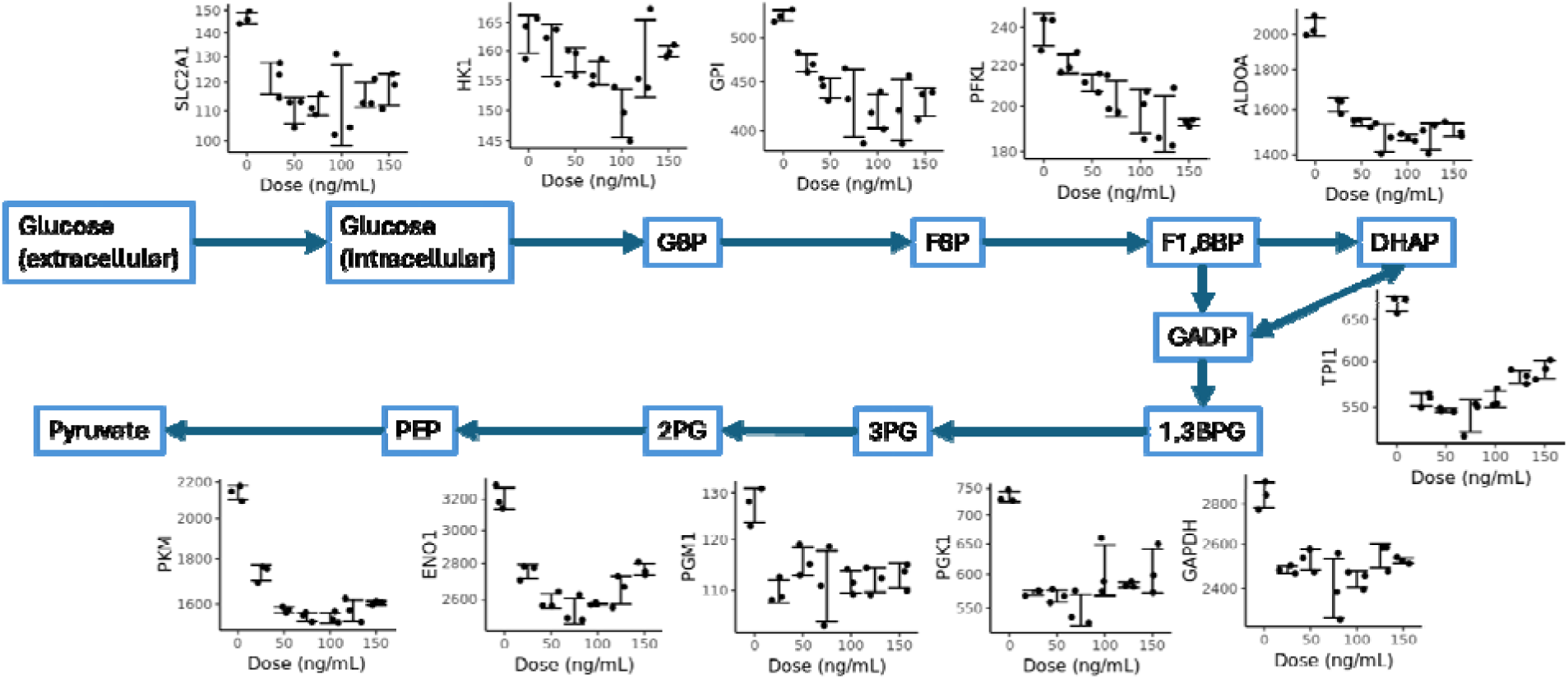
Canonical glycolysis genes were nonlinearly downregulated by increasing EtBr dosage. Read counts at 48 hours decreased initially but trended upwards at higher concentrations.

### 3.7 Variation in mtDNA-CN changes methylation of nuclear DNA CpGs nearby differentially expressed genes

Studies have shown that mtDNA-CN can alter nuclear DNA methylation sites and mtDNA-CN associated CpG sites can influence gene expression.^7^ We therefore investigated nuclear DNA methylation following EtBr treatment for matched samples. Specifically, to identify differentially methylated CpGs associated with mtDNA-CN reduction, we regressed genome-wide nuclear DNA methylation sites against mtDNA-CN. Among 786,095 tested CpG sites, 435 were significantly differentially methylated after multiple test correction (Fig. 5A), of which 234 CpGs were hypermethylated and 201 CpGs were hypomethylated upon mtDNA-CN reduction (Fig. 5B). Additionally, two differentially methylated CpGs, cg26094004 (adj.p = 3.11×10^-6^) and cg26563141 (adj.p = 4.20×10^-2^), had previously been associated with mtDNA-CN variation.^7^

**Fig. 5.**
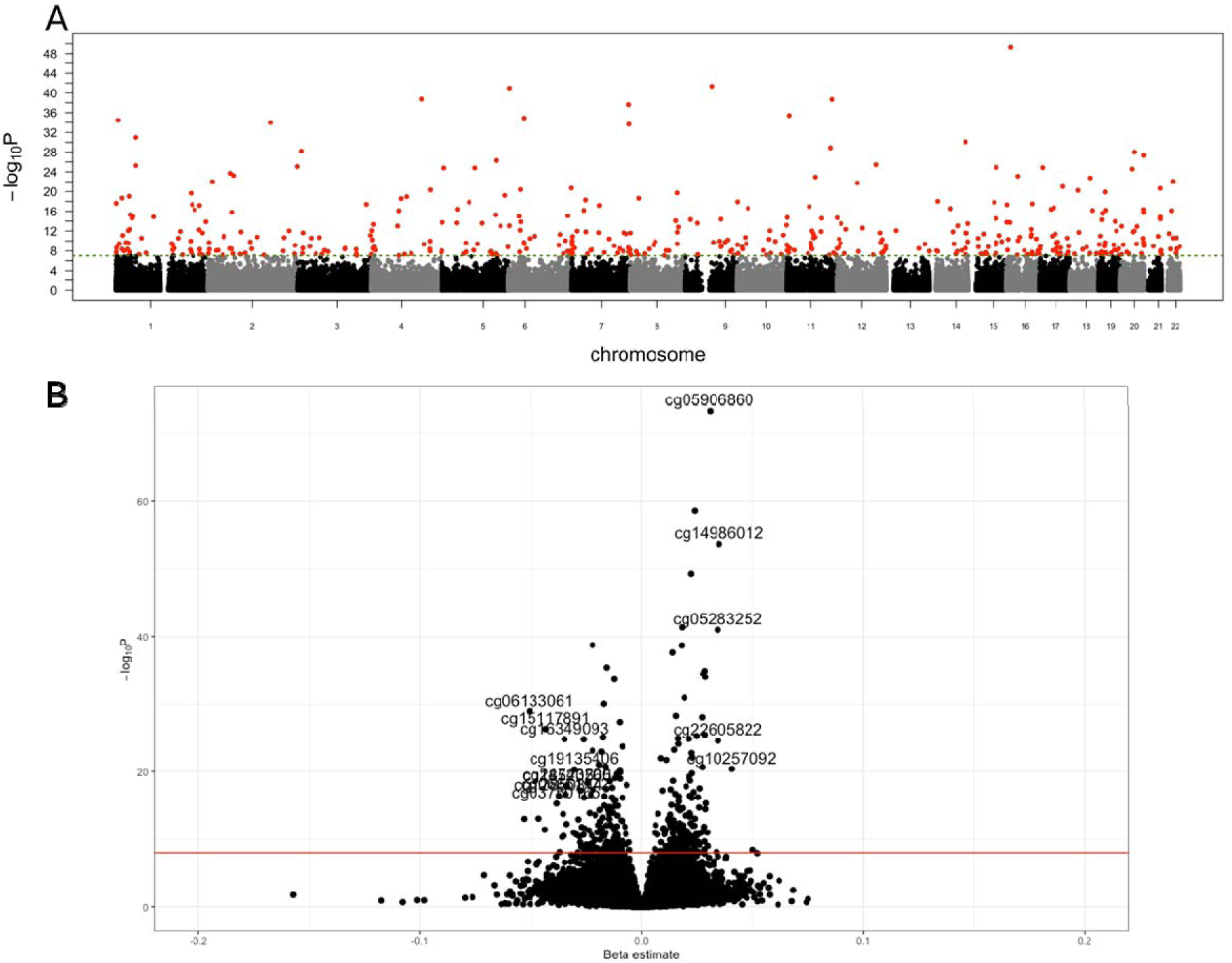
Ethidium Bromide treatment led to differential nuclear methylation. A) Linear regression of mtDNA-CN on methylation identified 435 CpGs with significant differential methylation. B) Volcano plot of methylation p-values and effect sizes.

Among the 390 differentially expressed genes, 229 genes had differentially methylated CpG sites located within 1 Mbp, which is 16% more than what would be expected by chance (P<0.023, 1000 simulations). Of the 164 unique differentially methylated CpGs located near differentially expressed genes, 97 were positively associated and 67 were negatively associated with reduction in mtDNA-CN, exhibiting a bias toward positively associated CpGs (Chi-square p = 0.019). Additionally, hypermethylated CpGs exhibited significantly larger effect sizes in methylation change compared to hypomethylated CpGs (Wilcoxon rank-sum test, p = 0.044). A CpG site of particular interest was cg26094004, which was associated with mtDNA-CN in both this study and a previous study.^7^ One of the DE genes within 1 Mbp of this CpG was the *ACLY* gene, an important regulator of cytosolic Acetyl-CoA and lipid biosynthesis.^42,43^ Gene ontology analysis of the 229 genes representing significant CpG-gene pairs identified overlapping GO terms found in the DE gene analysis, including protein binding (GO:0005515, adj.p = 2.29×10^-3^), membrane-bounded organelle (GO:0043227, adj.p = 2.02×10^-4^), small molecule binding (GO:0036094, adj.p = 3.71×10^-2^), and cytoplasm (GO:0005737, adj.p = 2.66×10^-4^). These gene ontology results from CpG-gene pairs suggests that mtDNA-CN induced differential methylation at these CpG sites regulates the expression of genes involved in cytosolic metabolic protein regulation through mitochondrial metabolites, such as *ACLY* produced acetyl-CoA.^44^

### 3.8 mtDNA-CN showed log-linear longitudinal response

Of the pathways and processes that we identified with dosage response, we wanted to distinguish upstream genes responding directly to mtDNA variation from downstream genes reacting to the cell state changes induced by the upstream genes. To assess the timing of cellular responses to EtBr treatment, we treated cells with 150 ng/ml EtBr for 48 hours, after which we removed EtBr, and allowed cells to recover for 144 hours. Samples were taken from experimental and control cells every 24 hours, except for hours 96 and 120. We observed that in treated cells, mtDNA-CN decreased in a log-linear fashion during the treatment phase, and mtDNA-CN increased with the same slope in the recovery phase (Fig. 6). At hour 144, recovering cells overcorrected and had mtDNA-CN higher than the controls. After hour 168, there were no significant differences in mtDNA-CN between the experimental and the control group.

**Fig. 6.**
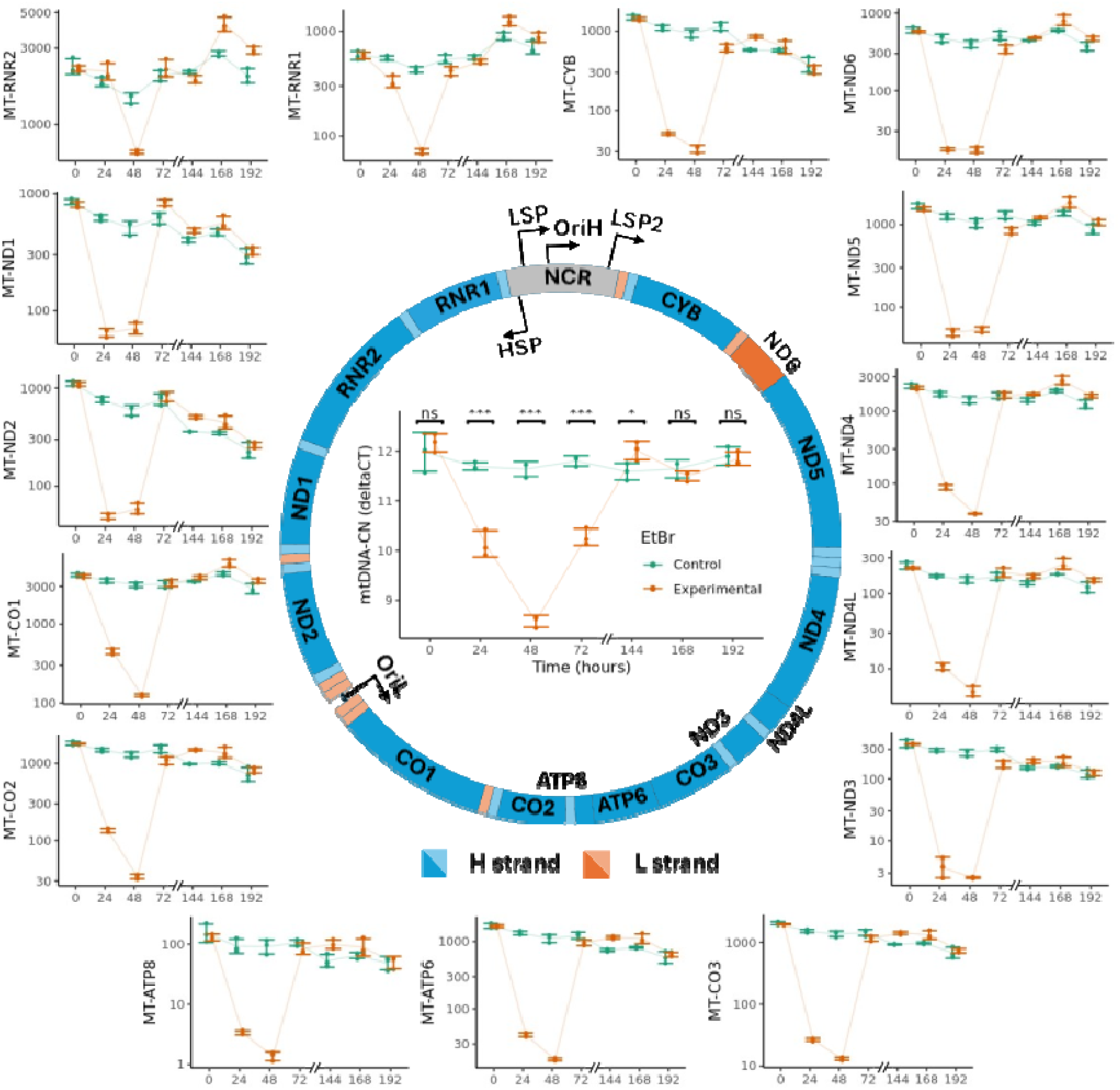
mtDNA-CN and mtRNA showed perturbation and recovery to EtBr introduction and removal. Plots showing changes in cellular mtDNA-CN and mtRNA counts as EtBr was introduced at hour 0 and removed at hour 48 (N=3). For comparison, measurements from cells grown in no EtBr are included. Error bars are +/-1SD.

### 3.9 Mitochondrially encoded genes show accelerated longitudinal response

Based on our observation that individual mtRNAs showed varied responses to EtBr treatment, we examined whether specific mtRNAs also exhibited different longitudinal changes (Fig. 6). While all mtRNAs had significantly decreased read counts at hour 48 before recovering to control levels at hour 72,three different patterns emerged for how the mtRNAs approached hour 48. The 2 ribosomal subunits showed very little change in read counts at hour 24, likely due to the increased stability of rRNA sequestered by ribosomal proteins.^40,45^ MtRNA for genes closer to the promoters (*MT-ND1, MT-ND2, MT-ND3, MT-ND5, MT-*ND6) showed similar read counts at hours 24 and 48, while genes further from the promoter (*MT-CO1, MT-CO2, MT-ATP8, MT-ATP6, MT-CO3, MT-ND4L, MT-ND4, MT-CYB*) showed higher read counts at hour 24 than at hour 48. MtRNAs respond to EtBr treatment and recovery faster than mtDNA-CN but showed positionally dependent variation reflective of the dosage response.

### 3.10 Differentially expressed genes show different longitudinal patterns in their treatment and recovery curves

To determine whether differentially expressed nuclear genes would exhibit varied longitudinal patterns, we calculated the logFC between treated and control groups at each time point and scaled the logFC to be zero at hour 0 and one at hour 48. We observed three longitudinal patterns among the top DE genes (Fig. 7): 1) log-linear behavior with scaled logFC from hours 0-72 of (0, 0.5, 1.0, 0.5); 2) switch-like behavior with scaled logFC (0, 1.0, 1.0, 0); and 3) temporally delayed behavior with scaled logFC (0, 0.25, 1.0, 1.0). It is interesting to note that the log-linear pattern resembles the longitudinal change in mtDNA-CN, the switch pattern resembles those of mtRNA, while the delayed pattern was unique to nuclear encoded genes.

**Fig. 7.**
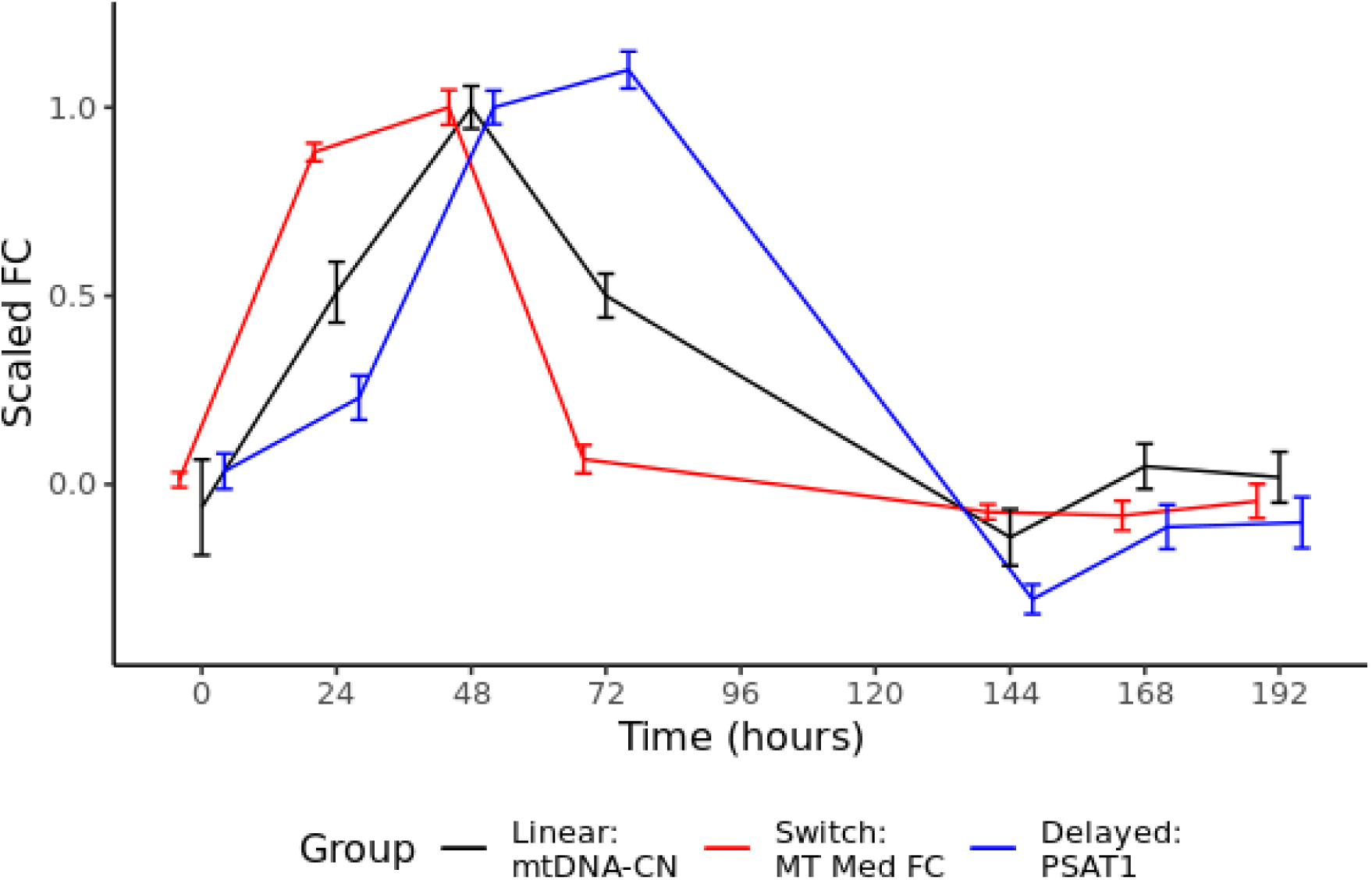
RNA counts and mtDNA-CN show different longitudinal treatment and recovery patterns. Schematic of longitudinal patterns for mtDNA-CN, median mtDNA gene, and a delayed nuclear gene (*PSAT1*). Y-axis shows logFC scaled to the maximum level of change at hour 48. Error bars are +-1SD.

For each differentially expressed gene, we calculated the Bayesian posterior likelihood for each pattern using the likelihood of observing the logFC at hours 24 and 72 given the expected logFC for that pattern. Prior likelihood was set at 1/3 for any of the patterns. A pattern was assigned to a gene if the likelihood for that pattern was at least twice that of the next likeliest pattern, and “ambiguous” if none of the patterns had a high likelihood (Supplementary Table 1).

Among the nuclear encoded DE genes, 109 showed a linear pattern, 49 showed a switch-like pattern, 107 had a delayed pattern, and 125 were ambiguous. Genes following the delayed pattern were significantly skewed towards being upregulated than the other categories (P = 3.70×10^-4^), although sensitivity analysis demonstrated that this imbalance was dependent on the posterior likelihood cutoff, with a more stringent cutoff making the delayed genes more skewed, while a relaxed cutoff lessens the imbalance (Table 2).

**Table 2.** Distribution of significant differentially regulated genes by the pattern of their treatment-recovery response. Sensitivity analysis is conducted by varying the required minimum ratio between the highest and second highest likelihood for a gene to be assigned a pattern. A minimum ratio of 2 is used to assign longitudinal patterns to genes.

| Pattern | Reg | Min ratio between 1st and 2nd |  |  |
| --- | --- | --- | --- | --- |
|  |  | 1 | 2 | 4 |
| Delayed | Up | 86 | 74 | 65 |
|  | Down | 61 | 33 | 29 |
| Linear | Up | 83 | 53 | 37 |
|  | Down | 86 | 56 | 40 |
| Switch | Up | 22 | 13 | 4 |
|  | Down | 52 | 36 | 26 |
| None | Up | - | 51 | 85 |
|  | Down | - | 74 | 104 |

### 3.11 Genes with delayed pattern were enriched for additional pathways

To test whether the narrowed focus for nuclear genes provided by the longitudinal patterns could provide additional insight into affected biological pathways, we performed overrepresentation analysis on the genes grouped by their longitudinal pattern and regulatory direction. This analysis revealed no significantly enriched GO biological processes for genes in the linear and switch pattern. Upregulated delayed genes were enriched for 42 terms, while downregulated delayed genes were enriched for 11 terms (Supplementary Fig. 2).

We compared enrichment results from before and after categorizing genes by their longitudinal pattern (Table 3, Supplementary Table 2). Comparing enrichment results from all upregulated genes and delayed upregulated genes, the 18 terms enriched by all upregulated genes were also enriched by delayed upregulated genes; in addition, delayed upregulated genes were enriched for response to nutrient levels (GO: 0031667, adj.p = 5.55×10^-4^). Comparing enrichment results from all downregulated genes and delayed downregulated genes, mRNA processing (GO:0006397, adj.p = 9.53×10^-3^) was shared by both but cholesterol metabolic process (GO:0008203, adj.p = 0.296) was not enriched by the delayed genes. New terms enriched by delayed downregulated genes included ribonucleoprotein complex biogenesis (GO:0022613, adj.p = 6.86×10^-3^) and response to fatty acid (GO:0070542, adj.p = 0.0130).

**Table 3.** Different GO terms were overrepresent by different longitudinal categories of differentially expressed genes. Within the table are the number of genes in each longitudinal category for each GO term. Cells highlighted in green indicate that the GO term was statistically significantly enriched by the genes in that category. In the left column, N indicates the total number of genes annotated under each GO term. In the bottom row, the number of nuclear encoded genes with valid entrez gene IDs under each category are displayed.

| Term | N | Upregulated |  |  |  |  | Downregulated |  |  |  |  |
| --- | --- | --- | --- | --- | --- | --- | --- | --- | --- | --- | --- |
|  |  | All | Delay | Linear | Switch | None | All | Delay | Linear | Switch | None |
| alpha-amino acid metabolic process | 123 | 11 | 8 | 1 | 1 | 1 | 3 | 1 | 1 | 1 | 0 |
| tRNA aminoacylation | 42 | 11 | 9 | 1 | 1 | 0 | 2 | 1 | 1 | 0 | 0 |
| amino acid import across plasma membrane | 29 | 6 | 5 | 0 | 0 | 1 | 0 | 0 | 0 | 0 | 0 |
| response to nutrient levels | 158 | 8 | 8 | 0 | 0 | 0 | 0 | 0 | 0 | 0 | 0 |
| mRNA processing | 422 | 3 | 1 | 2 | 0 | 0 | 21 | 8 | 6 | 1 | 6 |
| cholesterol metabolic process | 80 | 1 | 0 | 0 | 0 | 1 | 10 | 2 | 4 | 2 | 2 |
| ribonucleoprotein complex biogenesis | 444 | 5 | 0 | 4 | 0 | 1 | 18 | 9 | 4 | 0 | 5 |
| response to fatty acid | 27 | 1 | 1 | 0 | 0 | 0 | 3 | 3 | 0 | 0 | 0 |
|  |  | 191 | 74 | 53 | 13 | 51 | 199 | 33 | 56 | 36 | 74 |

### 3.12 Differentially methylated CpG sites nearby delayed pattern differentially expressed genes identify shared functions

Given that epigenetic regulation through DNA methylation can influence gene expression and is expected to display a delayed response stemming from methyltransferase catalytic and downstream RNA polymerase activity, we focused our downstream analysis on delayed pattern genes.^46^ We prioritized 107 DE genes that displayed a delayed pattern from the longitudinal study. Of the 107 genes, 45 of these genes were located within 1 Mbp of a differentially methylated CpG sites, which is 18% more than would be expected (P=0.018, 1000 simulations, Sup Table 2). As above, we employed Bayesian posteriors to identify longitudinal methylation patterns for these CpGs. Among these CpGs, one CpG site (cg18194483) exhibited a delayed longitudinal response, three CpGs (cg18194483, cg26312802, cg02404346) demonstrated linear longitudinal response, and 41 CpGs did not conform to a pattern (Supplementary Fig. 3).

Gene ontology analysis of the 45 genes representing significant CpG-gene pairs identified both unique and overlapping GO terms as compared to the DE gene dose response analysis. Unique terms found in the matched CpG-Gene analysis include cellular amino acid metabolic process (GO:0006520, adj.p = 3.17×10^-6^), peptide metabolic process (GO:0006518, adj.p = 3.10×10^-3^), cytoplasmic translation (GO:0006412, adj.p = 4.74×10^-3^), small molecule metabolic process (GO:0044281, adj.p = 1.26×10^-5^), and organic acid metabolic process (GO:0006082, adj.p = 6.95×10^-5^). This may suggest a distinct methylation mediated response focused on intracellular cytosolic metabolic function. Despite fewer genes in the CpG-gene pair GO analysis, several shared ontologies were identified in line with the DE results including tRNA aminoacylation (GO:0043039, adj.p = 1.26×10^-5^), amino acid import across plasma membrane (GO:0089718, adj.p = 1.04×10^-3^), and small molecule biosynthetic process (GO:0044283, adj.p = 4.49×10^-2^).) (Supplementary Fig. 4, Figure 3).

## 4 Discussion

Mitochondria are involved in multiple cellular pathways including cell signaling, biosynthesis, and bioenergetics.^47^ To determine what pathways are involved in mito-nuclear crosstalk, we perturbed mitochondrial function chemically and observed the changes in nuclear DNA methylation and gene expression. With increasing EtBr concentration and decreasing mtDNA-CN, we observed 390 differentially expressed nuclear encoded genes and 435 differentially methylated nuclear CpG sites. We also identified CpG–gene pairs consisting of significant genes within 1 Mbp of significant CpGs, with 229 differentially expressed genes mapped to 164 differentially methylated CpGs.

We observed that, with some exceptions, EtBr treatment decreased expression of mtDNA encoded subunits of the OXPHOS electron transport chain. Contrary to our expectations, the decrease in mtDNA and mtRNA abundance did not lead to an increase in expression for mitochondrial DNA replication and transcription genes, with equal numbers of genes being upregulated and downregulated in either pathway (Table 1). Conversely, oxidative phosphorylation had 14 upregulated genes and no downregulated genes, suggesting that the pathway was upregulated. These results further suggest that mito-nuclear cross talk is predominantly modulated by cellular energetics rather than mtDNA and mtRNA abundance, with the expression of mtDNA replication and transcription being tightly regulated.

While nuclear encoded mechanisms of mtDNA replication and transcription were not responding to mtDNA depletion, a closer examination of mtRNA levels showed dramatic responses within the mitochondria. Particularly conspicuous were the 4 genes immediately downstream of the HSP, which showed a nonlinear response to increasing EtBr concentration (Fig. 1). Since mtDNA replication is coupled to transcription, we hypothesized that the nonlinear response of these genes resulted from mtDNA depletion triggering an increase in transcription initiation, resulting in higher read counts at lower concentrations of EtBr. However, as EtBr concentration increases and mtDNA-CN decreases, the template for transcription was disappearing and increased initiation could no longer make-up for that shortfall, ultimately decreasing the number of reads mapped to these genes.

Increased transcription initiation did not necessarily correspond to increased transcription. While conventional dogma holds that mtDNA transcription covers the entire mitochondrial genome,^1,48^ suggesting that changes in mtRNA levels should be uniform, we instead observed uneven changes with genes further from the NCR showing greater logFC decrease than genes proximate to the NCR (Fig. 2). This suggests that EtBr treatment had decreased mitochondrial transcription processivity, with decreased read counts for genes further from the promoters explained by increased premature termination. The contradiction of increased initiation and decreased processivity could be explained by the dual role that transcription plays in the mitochondria, in that transcription is responsible for synthesizing both mRNA/rRNA and RNA primers for mtDNA replication. Transcription initiated from the LSP would terminate at the OriH to leave behind the RNA primer necessary for mtDNA replication, while full genome transcription would only occur if the polymerase continued uninterrupted past the OriH.^1,48^ Thus, we hypothesize that in EtBr treated cells, mtDNA replication was being promoted at the expense of mtDNA transcription.

While LSP has a canonical role in initiating mtDNA replication from OriH, we are unable to explain why it would be that the genes between HSP and OriL (*MT-RNR1, MT-RNR2, MT-ND1, MT-ND2*), showed strong nonlinear response to EtBr treatment. Several possible explanations present themselves. First, the LSP and HSP are located only 110 bp apart, with shared initiation mechanisms, suggesting that transcription initiation at both sites is coupled.^49^ In addition, the mechanism for early transcription termination for the LSP at OriH could be shared for the HSP at OriL. Second, increased transcription at the HSP may be intended to increase ribosomal RNAs levels, strengthening translational resources to compensate for overall decreased mtRNA abundance. Third, a noncanonical mechanism may exist for replication initiation at OriL starting from the HSP.

The pathways enriched among differentially expressed nuclear encoded genes included alpha-amino acid metabolic process, tRNA aminoacylation, amino acid import across plasma membrane, response to nutrient levels, ribonucleoprotein complex biogenesis, and cholesterol metabolic process (Fig 3, Table 3). In addition to cg26094004-*ACLY* CpG-gene pair, we identified three differentially methylated CpGs following a log-linear alteration pattern near the DE genes of *CTH*, *LSS*, and *SLC6A9*, which are involved in cysteine metabolism, sterol biosynthesis, and membrane protein transport, respectively.^50–52^ Gene ontology analysis of 45 significant CpG-gene pairs identified unique terms, cellular amino acid metabolic process, peptide metabolic process, cytoplasmic translation, small molecule metabolic process, and organic acid metabolic process suggesting a methylation-mediated response focused on intracellular cytosolic metabolic and translational function. Shared terms, such as tRNA aminoacylation, amino acid import, amino acid transmembrane transport, and small molecule biosynthetic process, aligned with DE genes results. Despite fewer genes in the CpG-Gene pair analysis, these shared pathways indicate that epigenetic changes further modulate the delayed transcriptional response to mtDNA-CN reduction, targeting amino acid metabolism, small molecule biosynthesis, and membrane transport. These results are replicated by other studies and indicate that cells undergoing mtDNA depletion undergo a change in cell state due to starvation response.^53–55^ Decreased aerobic respiration caused an increase in fatty acid oxidation, with cholesterol metabolic processes downregulated as free cholesterol levels increased.^56,57^ Meanwhile, other pathways were enriched as part of the cell’s stress response, increasing the availability of resources through upregulation of amino acid transport, metabolism, and aminoacylation, while downregulating energy intensive processes such as ribonucleoprotein complex biogenesis.

One pathway which we had expected to be upregulated was glycolysis, replacing OXPHOS as a major energy source for the cell.^41,58^ However, unlike work done in mtDNA depleted rho0 cells, our experiments were conducted before mtDNA was totally depleted in our cells, and measurements were taken while the cell state was still in flux. Thus, we observed glycolysis genes downregulated with decreasing mtDNA-CN (Fig. 4). Our results are supported by experiments showing decreased glycolysis in cells shortly after inhibition of the TCA cycle.^59^ We hypothesize that glycolysis is controlled by two opposing forces. On the one hand, decreased mtDNA-CN decreased OXPHOS and its corresponding consumption of pyruvate, causing a buildup of the glycolysis product and possibly leading to the downregulation of the pathway. On the other hand, decreased OXPHOS decreased ATP synthesis within the cell, and the energy shortage may lead to the replacement of OXPHOS by anaerobic respiration and upregulation of glycolysis. We hypothesize that forces favoring glycolysis would only overcome the downregulating factors if the energetic state of the cell was sufficiently desperate, either due to extended time spent with moderately decreased mtDNA-CN or shorter time with severe mtDNA depletion. This is supported by the nonlinear fishhook pattern we observed in some glycolysis genes, showing gene expression trending upwards for high EtBr concentration compared to intermediate EtBr concentration.

In summary, we conducted a study in HEK293T cell lines investigating the transcriptomic and methylomic effects of chemically induced mtDNA depletion. We found that the initial cellular response to decreased mtDNA-CN went against our expectations. Within the mitochondria, we observed a greater decrease in mtRNA than the decrease in mtDNA-CN, with increased production of mtDNA replication primers and decreased expression of genes. Among nuclear encoded genes, energy intensive processes were downregulated, and alternative energy production pathways were upregulated. In parallel, epigenetic regulation through differential CpG methylation near associated genes further modulated transcriptional responses. Furthermore, we were surprised to see glycolysis downregulated, suggesting the regulation of glycolysis was not solely dependent upon the energy needs of the cell. With this study, we elucidated mechanisms surrounding mtDNA replication and the initial response to mitochondrial dysfunction in the cell.

## CRediT authorship contribution statement

**Jiaqi Xie**: Conceptualization, Data curation, Formal analysis, Investigation, Methodology, Software, Visualization, Writing – original draft, Writing – review & editing**. Phyo W. Win**: Data curation, Formal analysis, Investigation, Methodology, Validation, Visualization, Writing – original draft, Writing – review & editing**. Charles Newcomb**: Data curation, Investigation, Writing – original draft**. Shaopeng Zeng**: Formal analysis, Visualization**. Christina A. Castellani**: Conceptualization, Funding acquisition, Investigation, Methodology, Project administration, Resources, Supervision, Writing – review & editing**. Dan E. Arking**: Conceptualization, Funding acquisition, Investigation, Methodology, Project administration, Resources, Supervision, Writing – review & editing.

## Funding

This study was made possible by funding from:

National Institutes of Health: R21AG082401 (DEA, CN, JX), R01HL170579 (DEA, CN, JX), R01AG085753 (DEA, CN, JX), R01HL144569 (DEA, CN, JX, CC).

Department of Pathology and Laboratory Medicine, Western University, Start-Up Funds to CAC, NSERC (Natural Sciences and Engineering Research Council of Canada), Grant Number RGPIN-2024-04933 to CAC, and Ontario Graduate Scholarship (OGS) to PWW.

## Appendix A. Supplementary data

Supplementary Tables. Supplementary Table 1, Supplementary Table 2

Supplementary Figures. Supplementary Figure 1, Supplementary Figure 2, Supplementary Figure 3, Supplementary Figure 4

