## Supplemental Figures for "Transcriptomic analysis of cells following decreased mitochondrial DNA-copy number reveals compensatory mechanisms in mitochondrial DNA replication and cellular energetics"

Supplementary Fig 1. Tree maps grouping similar enriched GO terms into categories. A) all upregulated genes. B) all downregulated genes. C) delayed upregulated genes. D) delayed downregulated genes.

A

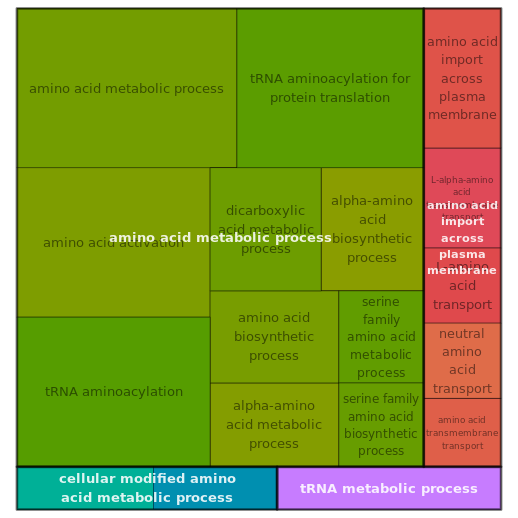

B

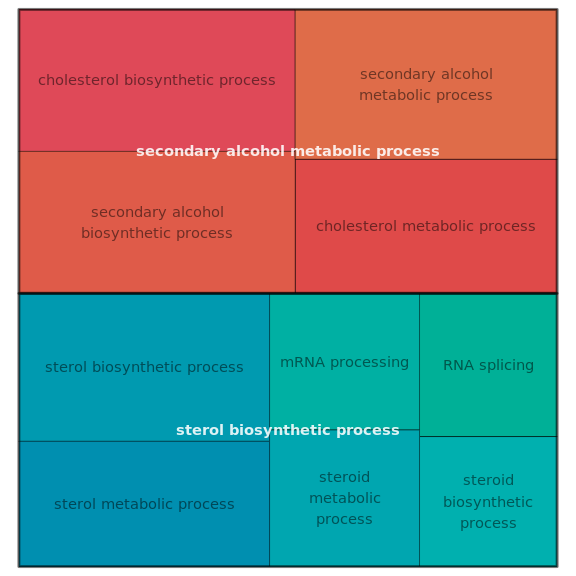

C

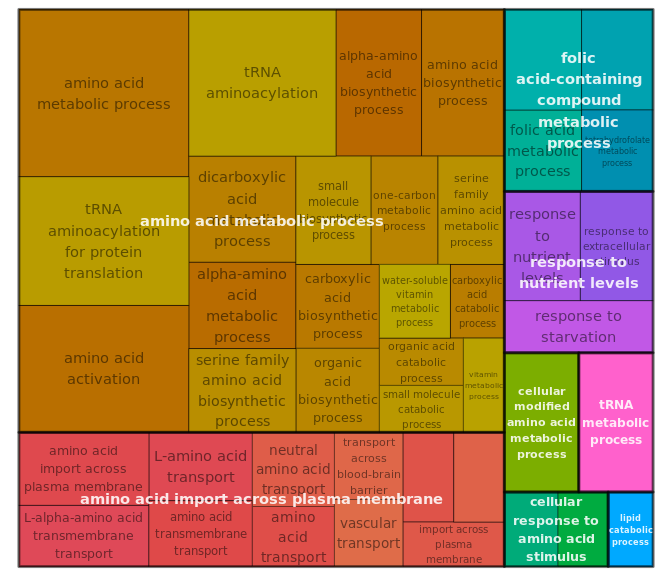

D

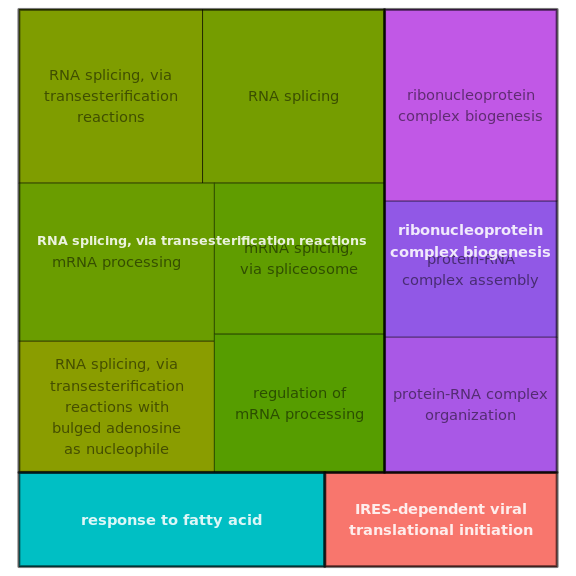

A

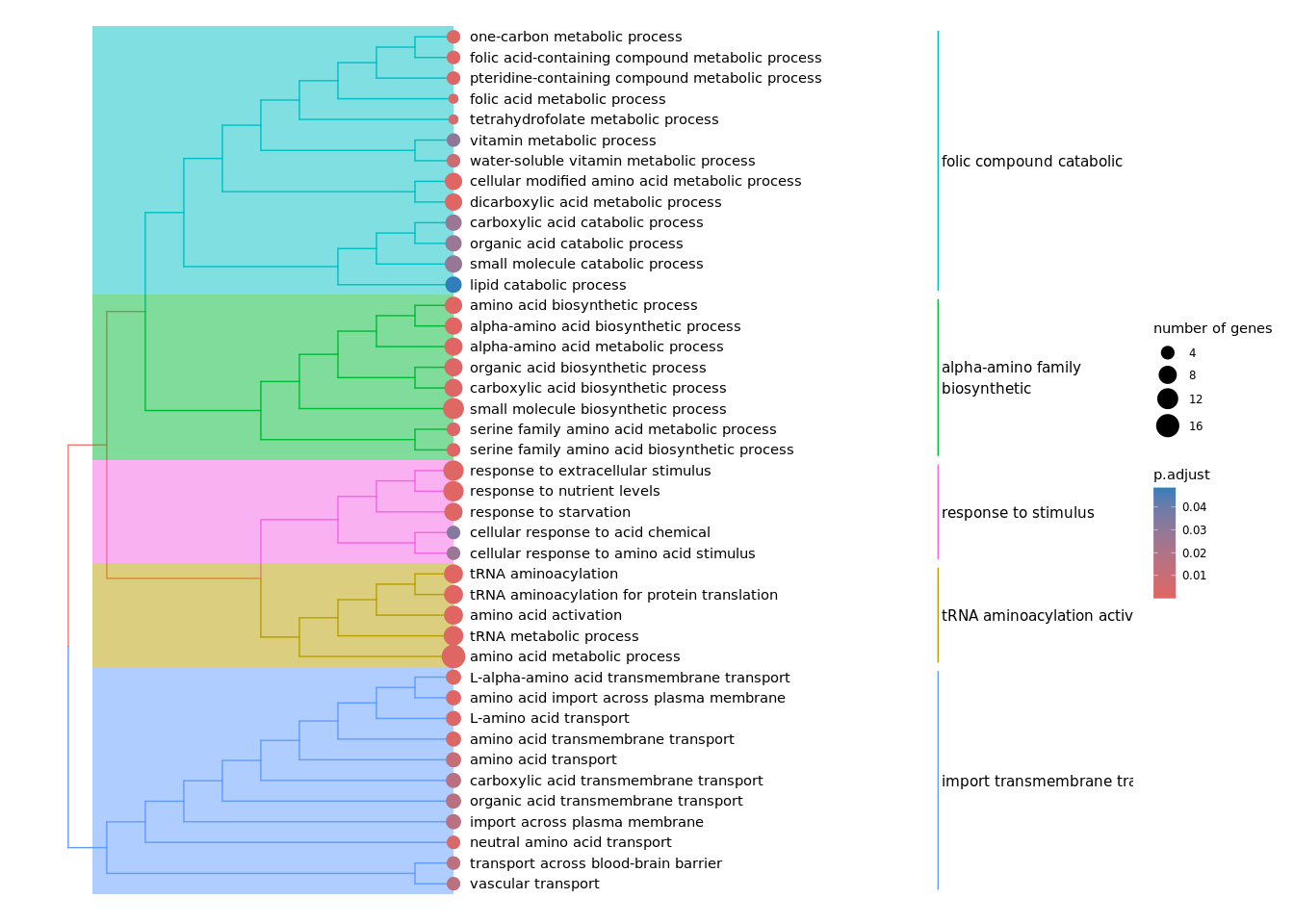

B

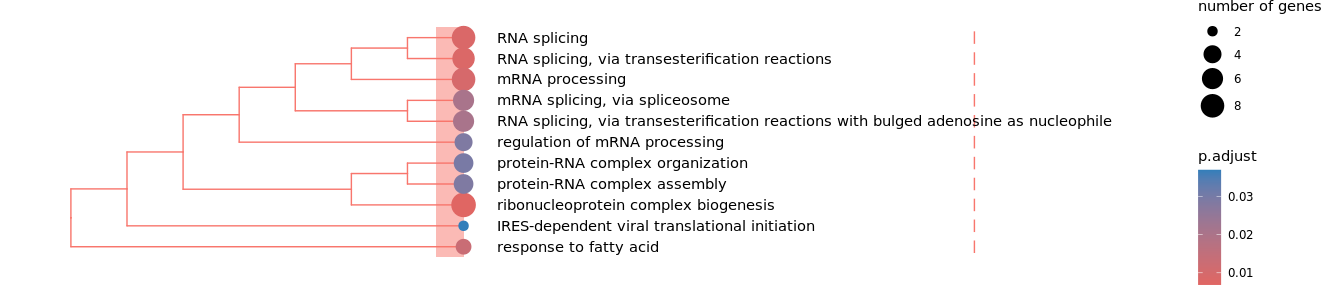

Supplementary Fig 2. Overrepresentation analysis of delayed pattern DE genes. Split here by A) upregulated and B) downregulated genes.

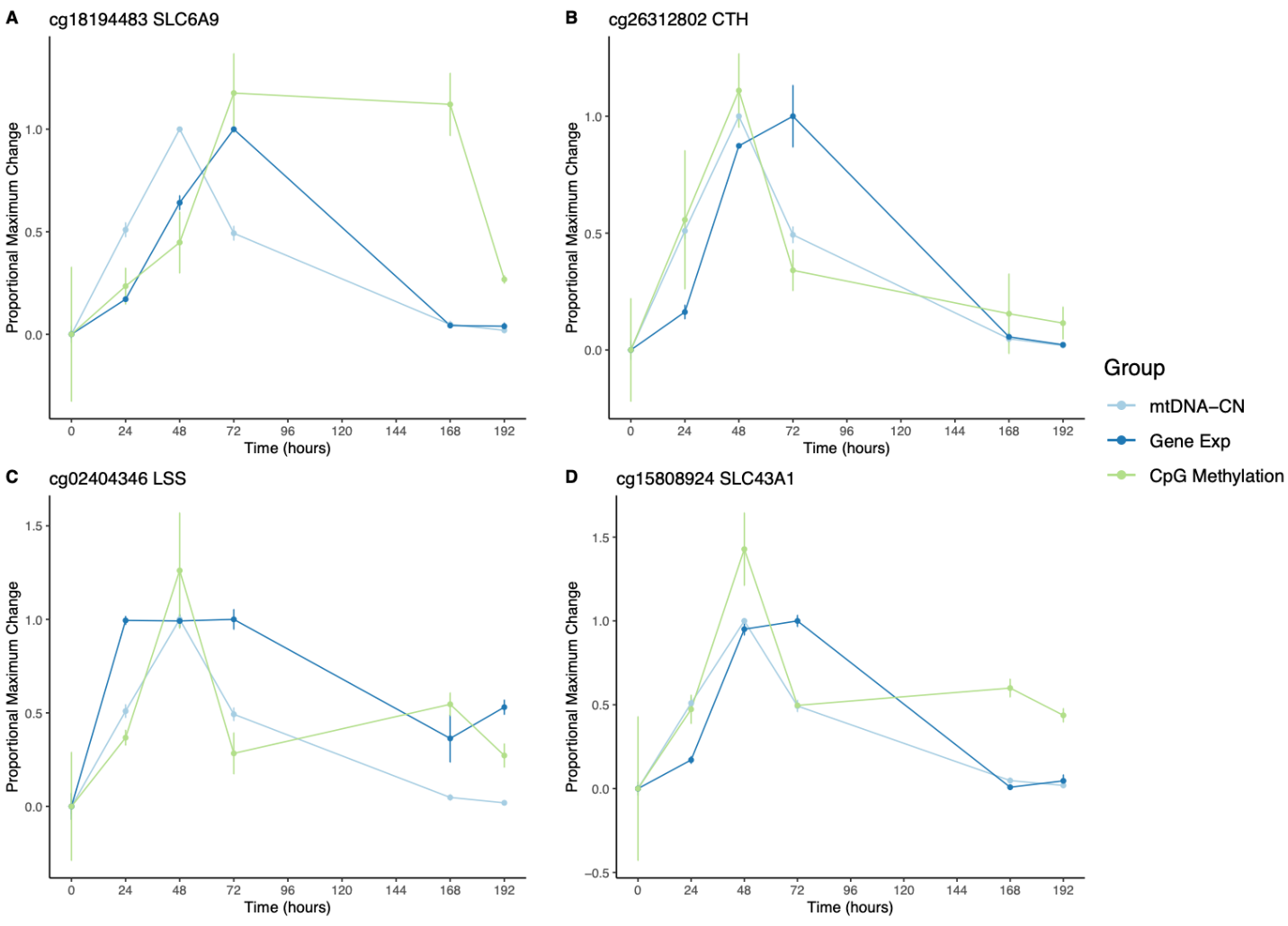

Supplementary Figure 3. Differentially methylated CpGs nearby differentially expressed delayed tracking genes display distinct longitudinal methylation patterns, A) one CpG sites exhibit a lagging pattern, with a delayed methylation response to the presence of EtBr. B-D) Three CpG sites show methylation change corresponding with mtDNA-CN change patterns.

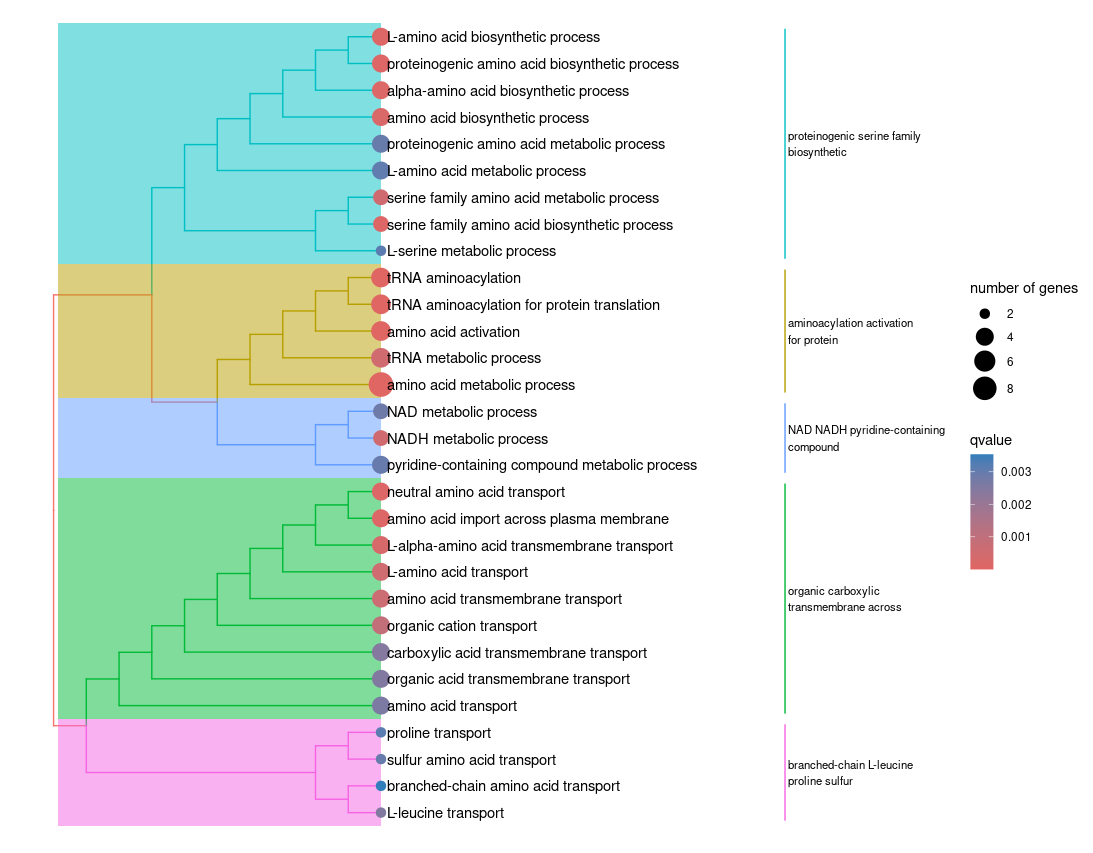

Supplementary Figure 4. GO Enrichment terms of differentially expressed delayed tracking genes nearby differentially methylated CpGs.
